# E2F1 Drives Breast Cancer Metastasis by Regulating the Target Gene FGF13 and Altering Cell Migration

**DOI:** 10.1101/671149

**Authors:** Daniel P. Hollern, Matthew R. Swiatnicki, Jonathan P. Rennhack, Sean A. Misek, Brooke C. Matson, Andrew McAuliff, Kathleen A. Gallo, Kathleen M. Caron, Eran R. Andrechek

**Affiliations:** Lineberger Comprehensive Cancer Center University of North Carolina; Department of Physiology, Michigan State University; University of North Carolina Department of Cell Biology

## Abstract

In prior work we demonstrated that loss of E2F transcription factors inhibits metastasis. Here we address the mechanisms for this phenotype and identify the E2F regulated genes that coordinate tumor cell metastasis. Transcriptomic profiling of E2F1 knockout tumors identified a role for E2F1 as a master regulator of a suite of pro-metastatic genes, but also uncovered E2F1 target genes with an unknown role in pulmonary metastasis. High expression of one of these genes, Fgf13, is associated with early human breast cancer metastasis in a clinical dataset. Together these data led to the hypothesis that Fgf13 is critical for breast cancer metastasis, and that upregulation of Fgf13 may partially explain how E2F1 promotes breast cancer metastasis. To test this hypothesis we ablated Fgf13 via CRISPR. Deletion of Fgf13 in a MMTV-PyMT breast cancer cell line reduces the frequency of pulmonary metastasis. In addition, loss of Fgf13 reduced *in vitro* cell migration, suggesting that Fgf13 may be critical for tumor cells to invade out of and escape the primary tumor. The significance of this work is twofold: we have both uncovered genomic features by which E2F1 regulates metastasis and we have identified new pro-metastatic functions for the E2F1 target gene Fgf13.

## INTRODUCTION

Breast cancer progression to metastatic disease is associated with poor prognosis, with only 22% of the patients surviving five years^54^. As a result, there is a critical need to understand the molecular mechanisms that regulate metastasis. High throughput transcriptomic assays have been pivotal in understanding alterations in the transcriptional programs of cancer cells during the various steps of metastasis. Repeated selection of cells with the propensity for organ-specific metastasis^7, 34, 47^ led to the discovery of transcriptional signatures unique to each metastatic site with characteristic transcriptomic changes^51^. Other studies have combined clinical observations with gene expression profiling to generate gene signatures that predict progression to metastasis^4, 68, 74^. Finally, gene expression profiling has been used to identify genes involved in metastasis which were then validated in genetically engineered mouse models of breast cancer^2, 11, 30, 71^.

One of the best characterized models of breast cancer metastasis is the MMTV-PyMT system where the expression of the polyoma virus middle T antigen is expressed under the control of the mouse mammary tumor virus promoter / enhancer^24^. Expression of the middle T antigen results in activation of key signaling pathways including Ras, PI3K/AKT and PLC-γ. These transgenic mice rapidly develop multifocal mammary tumors and develop pulmonary metastasis at endpoint with nearly 100% penetrance. Our recent work highlighted the similarities of MMTV-PyMT tumors to human breast cancer and identified shared gene expression alterations between this mouse model and human disease^28^. One example is our observation that E2F pathway signatures were elevated in the MMTV-PyMT model, and we ultimately validated the role for E2F1, E2F3, and E2F3 in tumor progression using mouse models^28, 30^. E2F transcription factors are canonically involved in the G1/S transition, ultimately either promoting (E2F1-3a) or suppressing (E2F3b-8) cell cycle progression^3, 13, 39^. In this study we expand upon this theme to demonstrate a role for the activator class of E2Fs in tumor progression independent of their role in cell cycle progression.

Interbreeding MMTV-PyMT mice with mice null for E2F1^16^, E2F2^49^ and E2F3^32^ resulted in alterations in mammary gland development, tumor latency, histology, and vascularization^29, 66^. In addition to, or perhaps as an effect of, the role of E2F1 and E2F2 on these tumor phenotypes, E2F1 and E2F2 deletion reduced metastatic capacity accompanied by a decrease in circulating tumor cells. This led to the hypothesis that the E2F transcription factors regulate tumor cell intrinsic gene expression programs that are critical for metastatic progression.

Given that E2F transcription factors have been demonstrated to bind thousands of individual target genes^5^, we sought to characterize the gene expression profiles of E2F1^−/−^ MMTV-PyMT tumors. We hypothesized that this approach would allow us to identify E2F1-mediated transcriptional programs which contribute to metastasis. This would allow us to identify E2F1 target genes which were previously implicated in breast cancer metastasis but would also allow to identify novel metastasis driver genes. In line with this, we have identified a suite of E2F1 target genes which have been previously implicated in breast cancer metastasis, suggesting that E2F1 may be a master regulator of breast cancer metastasis. In addition, this analysis has also identified a new role for the E2F1 target gene fibroblast growth factor 13 (Fgf13) in breast cancer metastasis.

## Results

### Genomic Comparison of E2F^WT/WT^ and E2F^−/−^ Tumors

To determine the global gene expression response to E2F loss, we analyzed MMTV-PyMT tumors from E2F^WT/WT^, E2F1^−/−^, E2F2^−/−^, and E2F3^+/−^ backgrounds on Affymetrix microarrays. Using an unsupervised classed discovery approach^73^, we investigated the gene expression relationships amongst the various tumors. We used 1000 iterations of data resampling to measure the frequency of co-clustering across 2-10 clusters. Examining potential classes using empirical cumulative distribution functions (CDF) showed that maximum CDF was reached with four clusters (Figure 1A). This suggests that this cohort of tumors can be divided into 4 distinct clusters, and adding additional clusters will have no statistically significant value^73^. To measure the correlation between samples within each cluster, we used silhouette width^58^ (Figure 1B). Silhouette width demonstrated the strongest correlations were present in cluster 1. In addition, we observe that the majority of samples have strong similarities to other samples within their assigned cluster. Sample co-clustering across the iterations of resampling is also illustrated in Figure 1C (see blue-white heatmap, where white shows 0% co-clustering to dark blue 100% clustering). Collectively, this analysis suggests the relationship of these tumors is best described by four clusters.

**Figure 1.**
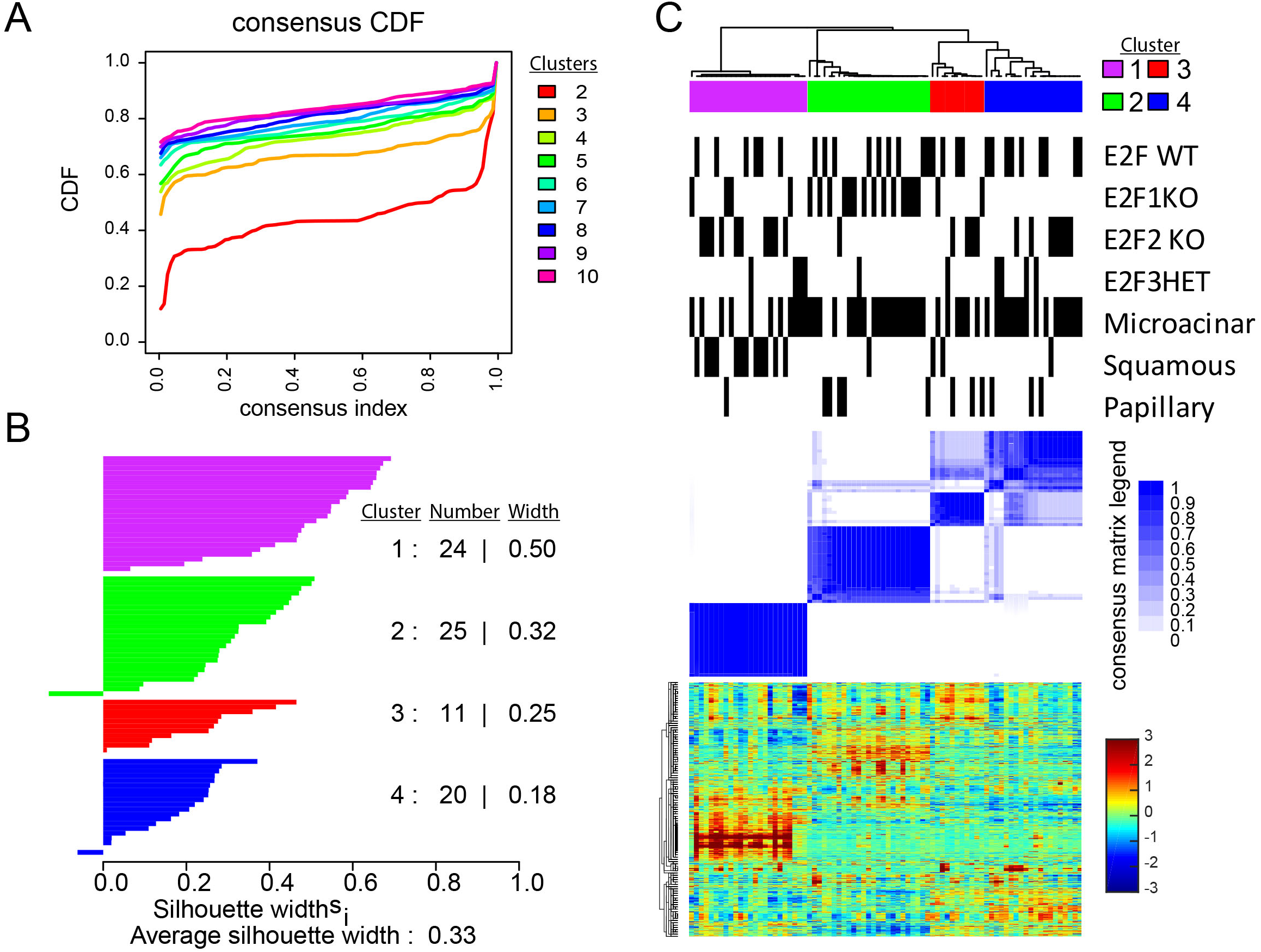
Consensus clustering reveals the major gene expression classes of MMTV-PyMT E2F knockout tumors. (A) Unsupervised class discovery of MMTV-PyMT tumor gene expression data (GSE 104397) by measure of empirical cumulative distribution functions^73^. The x-axis displays the consensus index, which is a measure of samples clustering together (0.0 samples that never cluster together, 1.0 samples that always cluster together). On the y-axis the CDF value is displayed. This provides a measure of cluster stability. Thus, this plot measures which number of clusters (as color coded) that provide maximum CDF (stable classification) for samples that are ambiguous (may not always cluster together). (B) A silhouette width^58^ analysis based upon the four clusters identified measures the correlation of samples within each cluster (the higher the silhouette width, the higher the correlation of a given sample to other samples in the cluster; each bar represent a sample). (C) The consensus cluster based upon the four selected clusters (as determined in A). Consensus clustering was performed using 1,000 iterations and 90% item (sample) resampling. The dendrogram across the top shows the relationship of samples on the basis of gene expression profiles. Below the dendrogram, color coded boxes itemize the cluster labels and concordant to those shown in panel B. Next, black bars provide sample annotations for genotype and tumor histology according to sample position in the dendrogram above and heatmaps below. Next, the blue heatmap illustrates the sample and cluster relationships over the 1,000 iterations of 90% resampling and cluster classification. The darkest blue indicates samples that co-clustered 100% of the time, white indicates samples that never clustered together. For samples with intermediate values, the percentage of co-clustering with other samples outside the cluster is shown. For example, the far left samples in cluster 2, sometimes clustered with samples in cluster 3 over the iterations of resampling a small percentage of the time. Alternatively, cluster 4 samples clustered with cluster 3 samples a small percentage of the iterations resampling and re-clustering. Finally, the heatmap shows the gene expression profiles of each sample (column-wise= samples, row-wise = genes). Genes were ordered using centroid linkage. Expression levels are shown according to the color-bar to the right of the heatmap. (Entire analysis) Data preprocessing: see methods.

Examining these clusters in more detail (Figure 1C), we observe that gene expression patterns correlate with both tumor histology and genotype. For example, cluster 1 (purple) featured mainly tumors with squamous histology. The microacinar and papillary tumors appeared to separate by genotype; with the majority of E2F1^−/−^ tumors ordered into cluster 2, E2F2^−/−^ and E2F3^+/−^ tumors in clusters 3 and 4, and E2F WT tumors were present in each cluster. As an additional analysis, we also used a supervised approach using a published gene set for intrinsic classification of mouse mammary tumors^53^. To adjust for centering biases^83^ and to enable accurate interpretation of this intrinsic analysis, we combined our dataset with published data^1, 2, 27^ using Bayesian Factor Regression Modelling^10^ (BFRM) to correct batch effects. This intrinsic analysis again separated tumors from the MMTV-PyMT model across distinct clusters (Figure S1A) and separated tumors according to histology and genotype (Figure S1B). In agreement with previous work^31^, the squamous tumors showed basal-like gene expression features (Figure S1B, i). MMTV-PyMT E2F1^−/−^ tumors clustered separately from other MMTV-PyMT tumors and showed high expression of some genes from the luminal cluster (Figure S1B, ii). As expected, we did not detect evidence for claudin low tumors across any MMTV-PyMT tumors (Figure S1B, iv). We did not observe differences in the proliferation cluster genes with regards to E2F status (Figure S1B, iii). However, we did observe a difference by tumor histological subtype. Observing median expression of the proliferation signature genes^15^, microacinar and squamous tumors showed high expression, while papillary tumors had significantly lower median expression (p<.01 papillary vs microacinar, or papillary vs squamous; Figure S2A). In agreement, retrospective analysis of tumor growth rate data, showed that papillary tumors progressed slower (days until 2,000 mm^3^) than microacinar or squamous tumors (Figure S2B). Taken together, these data demonstrate unique features of tumor histologies and tumor genotypes; with E2F1^−/−^ tumors exhibiting key molecular differences to other MMTV-PyMT tumors.

To test if these gene expression differences in E2F1^−/−^ tumors corresponded to differences in activation of major cell signaling pathways, we utilized a binary regression approach^6^ to predict pathway activation across the MMTV-PyMT tumors. This revealed E2F1^−/−^ tumors tend to have high activity of E2F4 (Figure 2A), and p53 (Figure 2B) pathways and low activity of pathways previously implicated in metastasis; the RhoA^78^, Src^50^, and Egfr^80^ signaling pathways (Figure 2C-E, respectively). Finally, using single sample gene set enrichment analysis (ssGSEA), we found that E2F1^−/−^ tumors had significantly lower expression of the Hallmark Hypoxia^41^ signature (Figure 2F); a process also associated with metastasis^42^.

**Figure 2.**
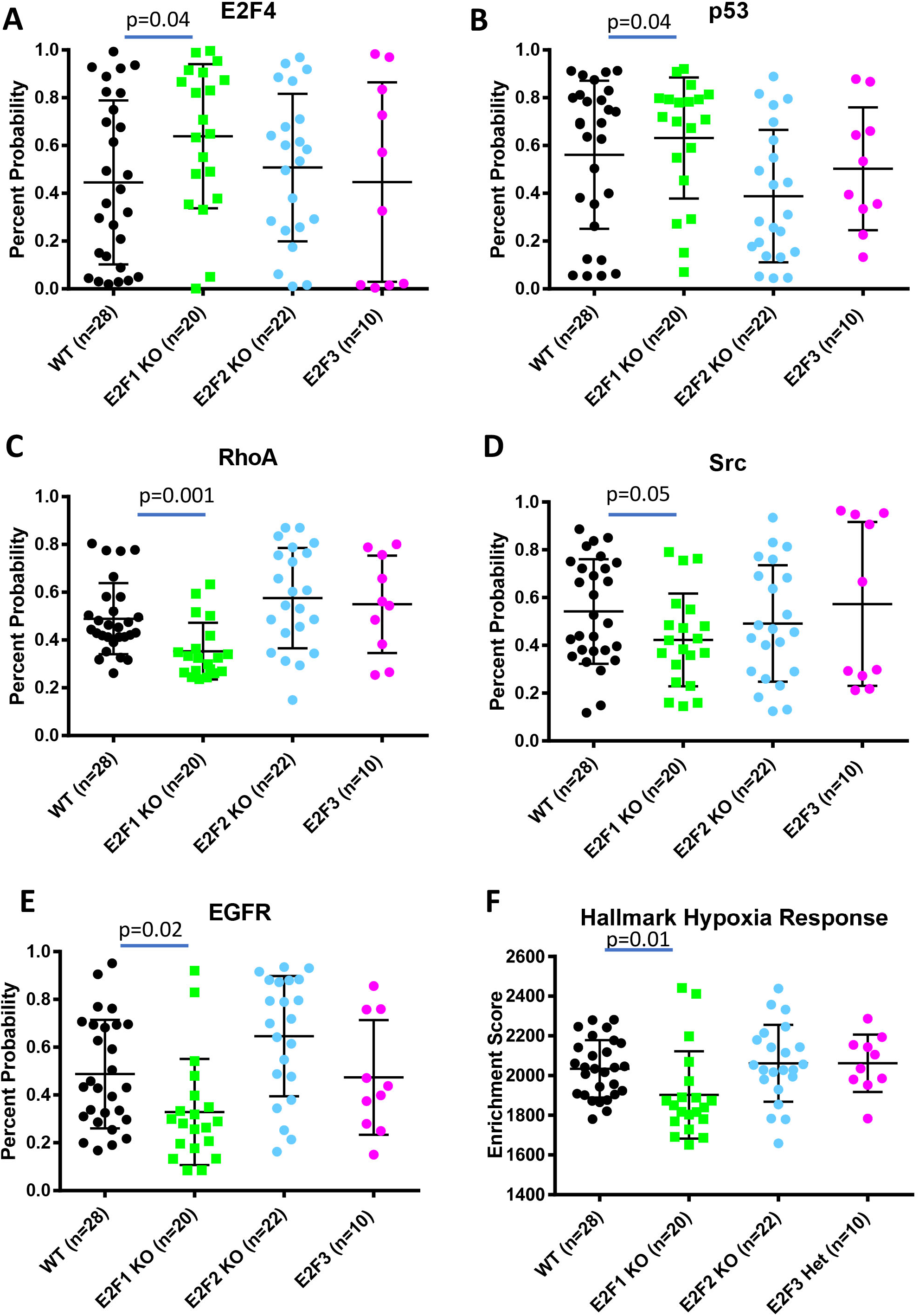
Gene expression signatures reveal pathway activation differences with loss of E2F1. (A) Using a binary regression approach^6, 28^ to predict E2F4 activity reveals significantly higher probability of E2F4 pathway activation in E2F1 KO tumors compared to E2F WT tumors (p=0.04). (B) Using a binary regression approach^6, 28^ to predict p53 activity reveals significantly higher probability of p53 pathway activation in E2F1 KO tumors compared to E2F WT tumors (p=0.04). (C) Using a binary regression approach^6, 28^ to predict RhoA activity reveals a significant reduction of probable RhoA pathway activation in E2F1 KO tumors compared to E2F WT tumors (p=0.001). (D) Using a binary regression approach^6, 28^ to predict Src activity reveals a reduction of probable Src pathway activation in E2F1 KO tumors compared to E2F WT tumors (p=0.05). (E) Using a binary regression approach^6, 28^ to predict Egfr activity reveals a significant reduction of probable Egfr pathway activation in E2F1 KO tumors compared to E2F WT tumors (p=0.02). (F) Using single sample gene set enrichment^41, 63^ to analyze expression of the Hallmark Hypoxia Response signature shows a significantly lower enrichment score in E2F1 KO tumors compared to E2F WT tumors (p=0.01).

To test for the genes that were significantly differentially regulated between E2F1^−/−^ and E2F^WT/WT^ tumors we used a supervised analysis^67^. Since we hypothesized that E2F1’s role in regulating metastasis was by transcriptional activation of target genes, we were particularly interested in the 226 genes that were significantly downregulated in the E2F1^−/−^ tumors. Many of the genes with low expression in E2F1 KO tumors were associated with hypoxia response (Supplemental file 1).

To begin characterizing these genes for metastatic potential, we used Kaplan Meier analysis of clinically and intrinsically annotated human breast cancer gene expression data^25^. To identify E2F1 target genes, we used ChIPBase^77^. Out of the 226 differentially regulated genes, 98 were E2F1 targets (supplemental file 1). From these, we focused on genes where high expression in tumors correlated with a decreased time to distant metastasis across all breast cancers as well as within with individual intrinsic subtypes of tumors. We identified 55 genes with pro-metastatic predictions (without discordant predictions in differing subtypes) (supplemental file 1); 34 of which had demonstrated E2F1 binding sites^5^. Using a Fisher’s exact test, we observed that the distribution of direct E2F1 targets was significantly higher in the genes concordant with human breast cancer metastasis predictions (p=0.001) than genes either discordant or not predictive of human breast cancer metastasis. This suggested that the E2F1 target genes altered in these tumors are associated with human breast cancer metastatic potential and highlights that E2F1 may be a master regulator of breast cancer metastasis since it controls the expression of multiple pro-metastatic genes.

The 55 genes identified above that were associated with human breast cancer metastasis were examined in the literature to identify which of the molecular changes have already been demonstrated to regulate breast cancer metastasis *in vivo*. As depicted in Figure S3, Vegfa^59^, Hbegf^84^, Hspb1^23^, Flt1^65^, L1cam^82^, and Plaur^76^ had significantly lower expression (p<0.05) in E2F1^−/−^ tumors and have all previously been shown to regulate breast cancer metastasis *in vivo.* Additionally, there were genes with significantly lower expression in E2F1^−/−^ tumors that had been shown to have *in vitro* invasion or migration function such as Areg^26^, Tead1^81^, Coro1C^69^, Lama5^22^, Tgm2^45^, and Fgf 7^79^ (p<0.05, Figure S4). Taken together, this shows that E2F1^−/−^ tumors have low expression of genes and pathways demonstrated to promote breast cancer metastasis.

### Testing additional genes for metastatic function

Since we had identified a number of pathways and genes altered in E2F1^−/−^ tumors which were previously identified as metastasis driver genes, we sought to identify novel regulators of metastasis from the genes downregulated in E2F1^−/−^ tumors. Candidate genes for further testing were prioritized by selecting genes that had not previously been demonstrated to regulate breast cancer metastasis, that were E2F target genes, and correlated with decreased time to distant metastasis across several human breast cancer human breast cancer subtypes. With this pipeline we identified differential expression of fibroblast growth factor 13 (*Fgf13*). As shown in Figure 3A, expression was significantly reduced 1.4-fold in E2F1^−/−^ tumors (q=0.01, p=0.02). Analysis of the 500 bp sequence upstream of the transcriptional start site (TSS) revealed seven E2F binding motifs in human and ten in the mouse sequence for this gene. High expression of Fgf13 was significantly associated with faster onset of distant metastasis across all cases of breast cancer (Figure 3B) but was not significant across all Er+ tumors (Figure 3C). However, it was significantly associated with accelerated time to distant metastasis in Er-tumors (Figure 3D). Observing intrinsic subtypes, Fgf13 was not associated with metastasis of Luminal A tumors (Figure 3E). However, in Luminal B, Basal-like, and Her-2 enriched tumors high expression Fgf 13 was predictive of early onset metastasis (Figure 3F-H respectively).

**Figure 3.**
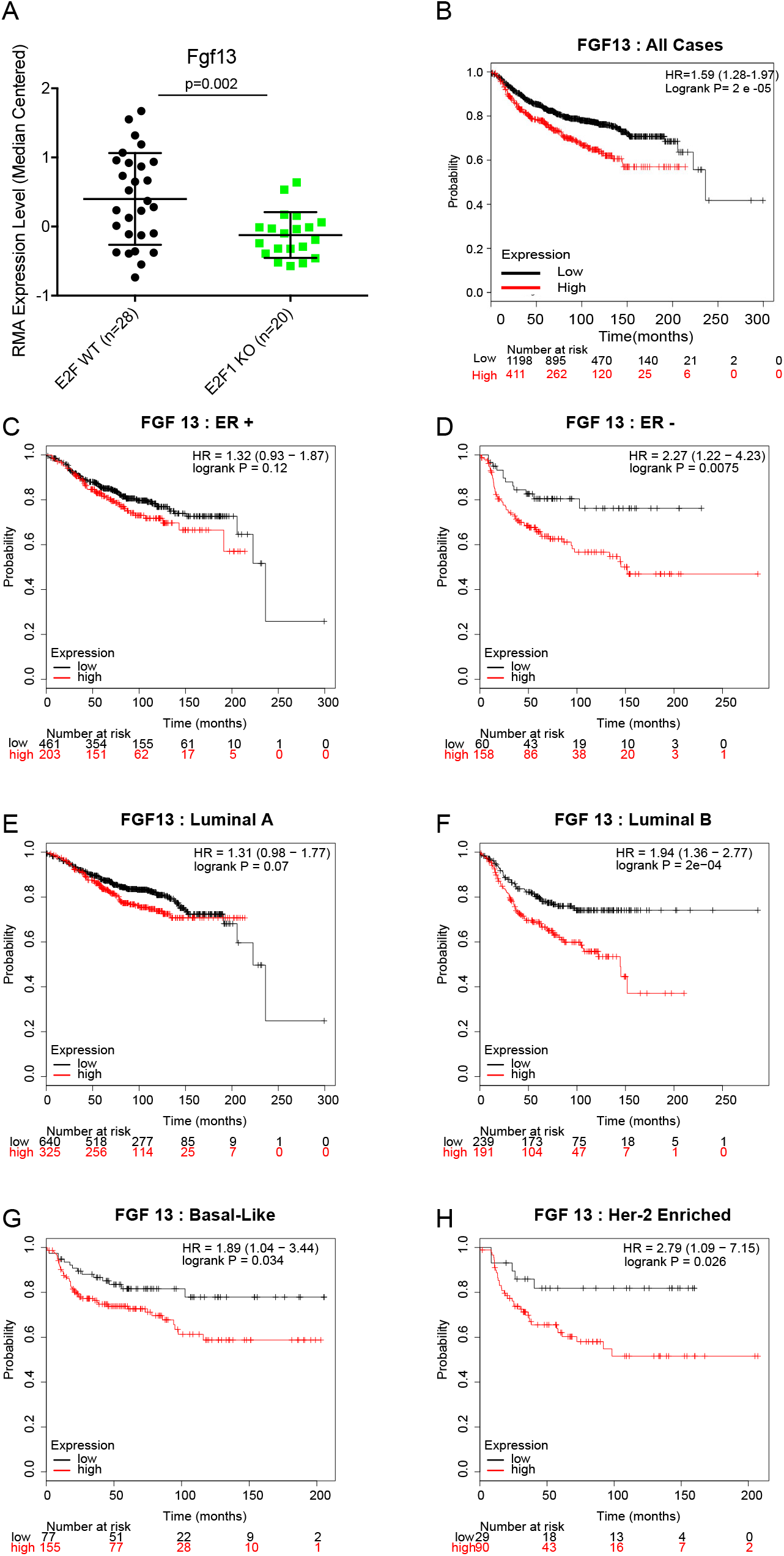
Expression of Fgf 13 is in E2F1 knockout tumors and association with human breast cancer metastasis. (A) Boxplot of RMA-normalized median centered expression levels of Fgf13 across MMTV-PyMT tumors shows a significant reduction of expression in E2F1 KO tumors (p=0.002, SAM: q=.01, 1.4 fold change). (B) Kaplan-Meier analysis of distant metastasis free survival across a dataset of human breast cancer^25^ shows that high expression of Fgf13 is associated with accelerated onset of metastatic progression (HR=1.59, logrank p= 2e^−05^). (C) Kaplan-Meier analysis of distant metastasis free survival across a dataset of human breast cancer^25^ shows that high expression of Fgf13 is associated (but not statistically significant) with accelerated onset of metastatic progression in estrogen receptor (ER) positive breast cancers. (D) Kaplan-Meier analysis of distant metastasis free survival across a dataset of human breast cancer^25^ shows that high expression of Fgf13 is associated with accelerated onset of metastatic progression in ER-negative breast cancers (HR=2.27, logrank p= 0.0075). (E) Kaplan-Meier analysis of distant metastasis free survival across a dataset of human breast cancer^25^ shows that high expression of Fgf13 is associated (but not statistically significant) with accelerated onset of metastatic progression in luminal A breast cancers. (F) Kaplan-Meier analysis of distant metastasis free survival across a dataset of human breast cancer^25^ shows that high expression of Fgf13 is associated with accelerated onset of metastatic progression in luminal B breast cancers (HR=1.94, logrank p=2e^−04^). (G) Kaplan-Meier analysis of distant metastasis free survival across a dataset of human breast cancer^25^ shows that high expression of Fgf13 is associated with accelerated onset of metastatic progression in basal-like breast cancers (HR=1.89, logrank p=0.034). (H) Kaplan-Meier analysis of distant metastasis free survival across a dataset of human breast cancer^25^ shows that high expression of Fgf13 is associated with accelerated onset of metastatic progression in Her2-enriched breast cancers (HR=2.79, logrank p=0.026).

To test this gene for metastatic behavior, we utilized a PyMT-derived cell line (PyMT 419 cells)^44^ and a CRISPR (clustered regularly interspaced short palindromic repeats) approach to create Fgf13 knockout cells. Figure 4A shows an example of sequence trace for Fgf13 control and a knockout lines. Western blot analysis confirmed that protein levels of Fgf13 were impacted by CRISPR gene editing (Figure 4B). In addition, we transduced our knockout lines with a vector to restore Fgf13 and confirmed re-addition by Fgf13 by western blot (Figure 4B).

**Figure 4.**
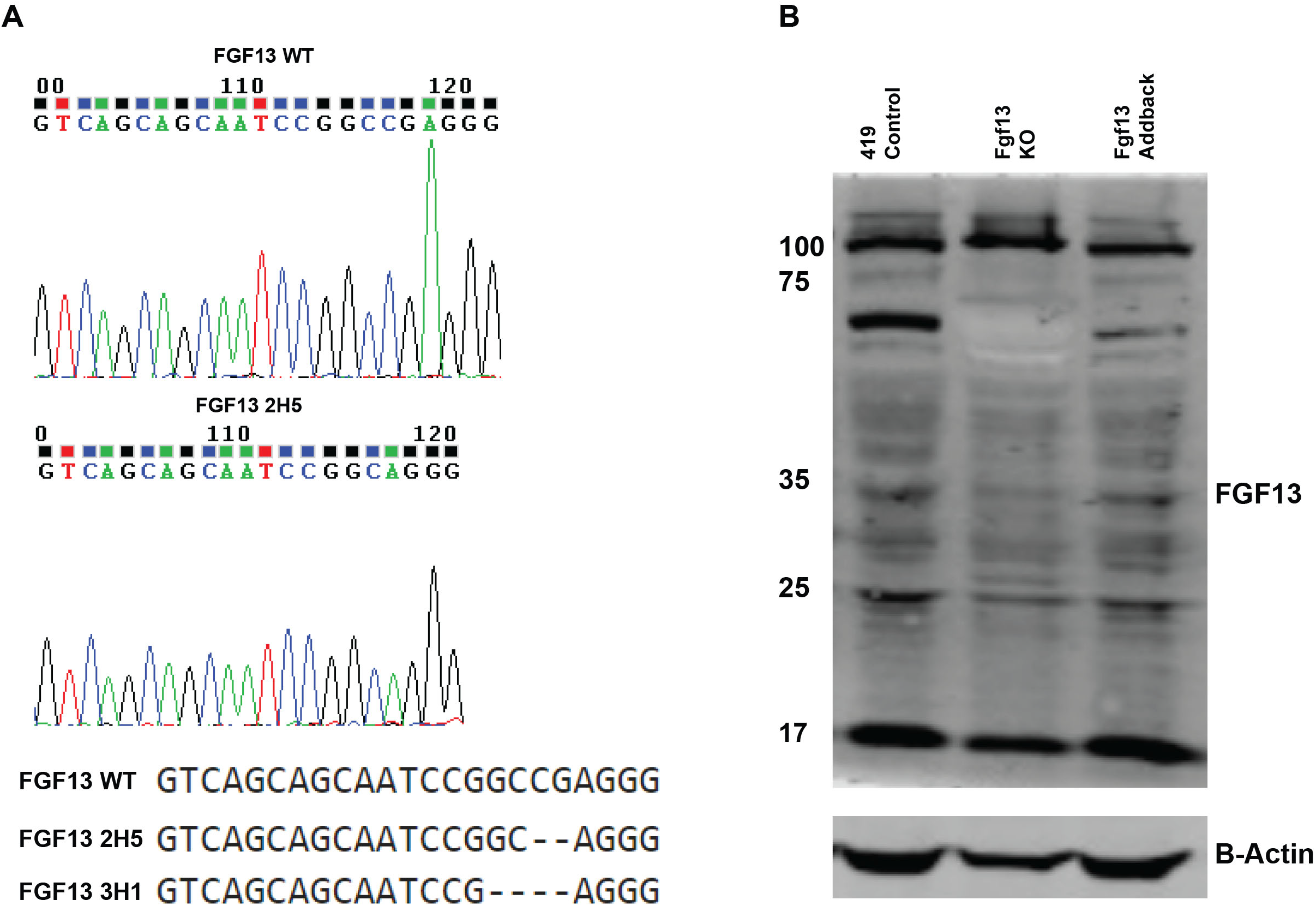
Sequence trace and alignment for CRISPR-mediated Fgf13 knockout. (A) Sequence trace for Fgf13 WT cells and Fgf 13 knockout clone 2H5. (B) Western blot analysis for Fgf13 WT cells, Fgf13 knockout cells, and Fgf13 add back cells.

We opted to assess metastatic capacity by tail vein injection. In this setting 50,000 cells were injected into the tail vein and lung colonization was examined 21 days later. In mice receiving control cells, robust metastatic colonization of the lungs was observed (Figure 5A). In addition, metastatic tumors were found sporadically throughout the mouse at other sites. Loss of Fgf13 dramatically limited the ability of the tumor cells to form metastases at the lung (Figure 5B) and re-expression of Fgf13 restored metastasis (Figure 5C). There was a significant reduction in the number of lesions present in the lungs for mice receiving Fgf13^−/−^ cells (Figure 5D). Together, these results show that Fgf13 regulates breast cancer pulmonary metastasis.

**Figure 5.**
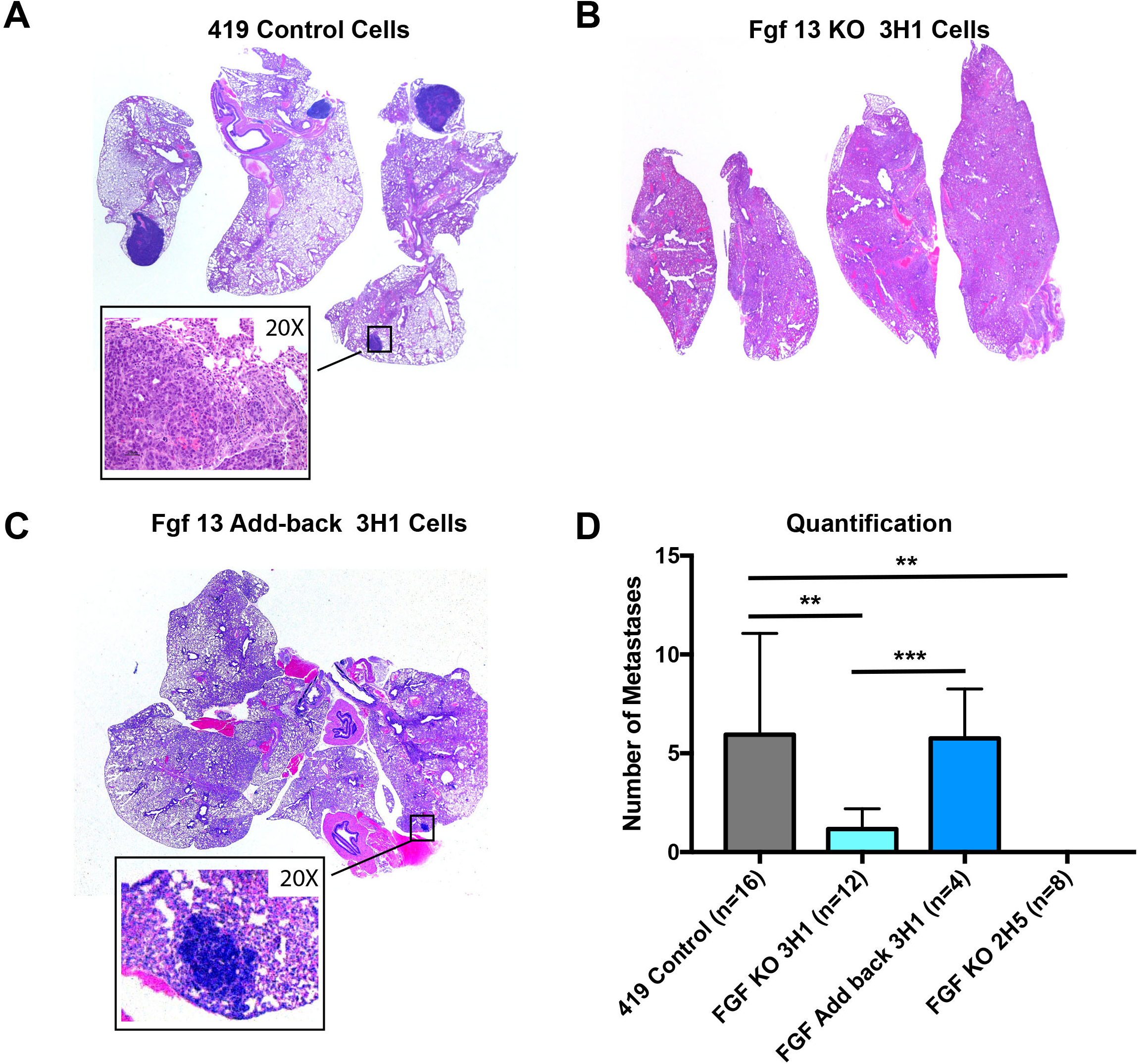
Fgf13 function in metastatic colonization of the lungs. (A)Representative image of lungs from mice receiving 419 control cells via a tail vein injection shows robust metastatic colonization of the lungs. (B) Lungs of mouse receiving the Fgf13 knockout clone 3H1 shows that loss of Fgf13 impaired tumor cell colonization of the lungs. (C) Lungs of mouse receiving the Fgf13 addback (to clone 3H1) shows that restoration of Fgf13 allows tumor cell colonization of the lungs. Quantification of FGF13 KO and addback clones compared to MMTV-PyMT 419 cell line control cells (black bar) shows a significant reduction in metastasis to the lungs in mice receiving or FGF13 (green bars) knockout cells (unpaired t-test, p<0.05; each clone compared by to wild type control).

### Investigating Fgf13 function

To predict Fgf13 function, we examined the Fgf13 covariance network using WGCNA analysis of primary MMTV-PyMT E2F wild type tumors (supplemental file 2), identifying 38 probes corresponding to 35 genes that tightly correlate with Fgf13 expression (gs threshold =0.6, p-value <0.0005). MSigDB revealed a significant association with a signature for genes up-regulated in invasive ductal carcinoma relative to ductal carcinoma in situ (Supplemental File 3, p-value =1.02 e^−4^, FDR q-value =3.97 e^−2^). We were also interested to note association with the Rac1 cell motility pathway (Supplemental File 2, p-value = 5.38 e^−7^, FDR q-value = 1.29 e^−3^). Key to this association was strong covariance with Rac1 (gs=0.60), the GTPase activating protein chimerin 1 (Chn1, gs=0.60), and Wasf1 (which acts downstream of Rac1 to regulate the cytoskeleton, gs=0.61). Matching this association, Fgf13 was part of the KEGG pathway for regulation of the actin cytoskeleton. Testing these alterations for the presence of an interaction network with Rb-E2F1 and the pathways altered in E2F1^−/−^ tumors illustrated a relationship between Fgf13, cytoskeleton and motility genes, and pathways predicted to have low activity in E2F1^−/−^ tumors (Figure 6A). WGCNA identified additional genes with cytoskeleton regulatory function such Tubulin beta 6 (Tubb6) and microtubule-associated protein 1B (Mtap1B). In addition, given Fgf13’s published role in neuronal cell differentiation^75^, we were interested to note this as another prominent theme amongst the Fgf13 metagene (Table 1). In testing these genes as a signature for association with human breast cancer metastasis events, we found that high expression of these genes was significantly associated with a shorter time to distant metastasis across breast cancer tumors (Figure 6B).

**TABLE 1:** DEVELOPMENTAL AND MIGRATION THEMES IN FGF13 METAGENE

| **Gene** | **Gene Significance Score** | **Function / Expression Pattern** | **Ref** |
| --- | --- | --- | --- |
| BEX4 | 0.72 | Family of Proteins involved in Neuronal Differentiation. | 9 |
| KCDN2 | 0.67 | Potassium ion channel protein expressed in neurons | 46 |
| Mtap1b | 0.67 | Stabilizes microtubules and implicated in neuronal migration | 48 |
| KLF16 | 0.64 | Regulation of neurite outgrowth | 70 |
| SEMA3<br>C | 0.61 | Regulation of neuronal migration | 72 |
| WASF1 | 0.61 | Neurite outgrowth | 12 |
| Rims2 | 0.6 | Important to neuron synapse function | 33 |
| Itm2a | 0.6 | Motor neuron marker | 61 |
| Rac1 | 0.6 | Neuronal cell migration | 35 |
| Chn1 | 0.6 | Regulation of neuron dendrite morphology | 8 |

**Figure 6.**
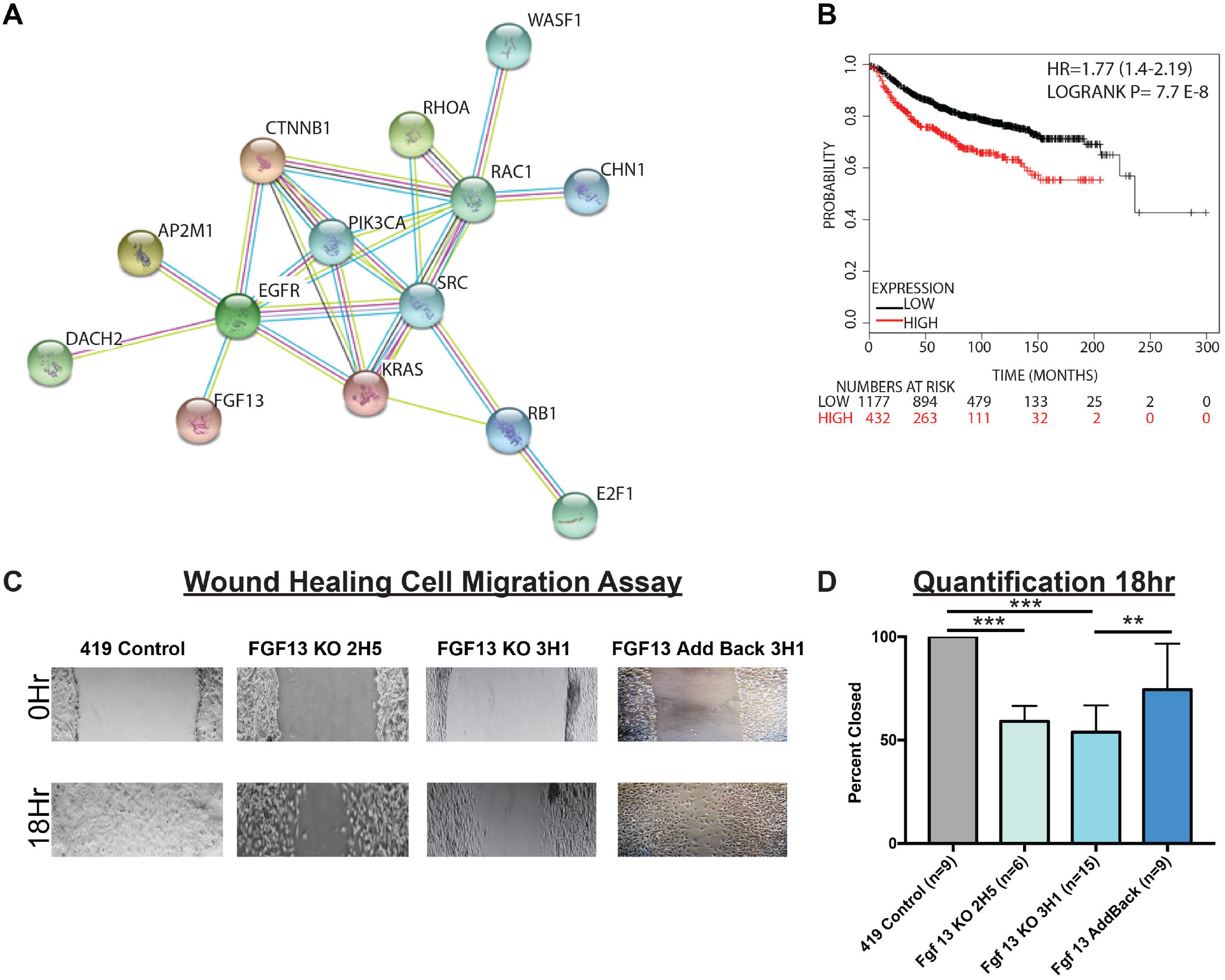
WGCNA analysis of MMTV-PyMT tumors predicts Fgf13 function in cell motility with confirmation by wound healing assay. (A) String interaction network for Fgf 13 covariance network genes and cell signaling pathways with low activity in E2F1-/- tumors reveals an association for genes with a gene significance score greater than 0.60 shows FGF13 many of the the FGF13 metagene are demonstrated interactors, including genes that function in cell motility (Rac1, Wasf1, Chn1). In addition, these covariance network genes associate with the Ras, Egfr, RhoA, Src, Rb, E2F1, and beta-catenin pathways. (B) Kaplan Meier analysis for the Fgf 13 covariance network genes as a signature shows these genes are significantly associated with earlier human breast cancer metastasis. (C) Scratch assay photos showing impaired migratory ability of 419 control cells, FGF13 KO cells, and Fgf13 add-back cells. (D) Quantification wound closure at 18 hours in of 419 control cells, FGF13 KO cells, and Fgf13 add-back cells.

To test for a possible function in cell motility, we assessed the migratory capacity of the MMTV-PyMT control cells and Fgf13 KO cells using a scratch assay (Figure 6C). While control cells were able to close the scratch at 18 hours, the Fgf13 KO clones demonstrated a significant defect in cell migration. Re-expression of Fgf13 restored migratory capacity of these cells, with wounds nearly closed by 18 hours (Figure 6D). Together, these data confirm the bioinformatic predictions that Fgf13 functions in cell migration and provides a likely explanation for the metastasis defects associated with Fgf13 loss.

## DISCUSSION

Using a transcriptomic approach, we investigated the mechanism by which E2Fs regulate breast cancer metastasis^30^. Here we used bioinformatic analysis to compare global gene expression differences between E2F^WT/WT^ MMTV-PyMT tumors and E2F1^−/−^ MMTV-PyMT tumors. We demonstrated that loss of E2F1 led to decreased activity in several key signaling pathways previously demonstrated to regulate metastasis (Figure 2). Many of the genes downregulated in E2F1^−/−^ tumors correlate with a faster progression to metastatic disease in human clinical data were significantly associated with the gene expression response to hypoxia (Supplemental File 1).

The majority of the genes associated with hypoxia were also direct E2F1 target genes. Hypoxia has been described as a master regulator of metastasis due to the result of gene expression changes brought about by hypoxia response^42^. These gene expression changes enable tumor cells to progress through multiple checkpoints during the metastatic cascade. This includes promoting angiogenesis, epithelial to mesenchymal transition, tumor cell invasion, remodeling of the extra cellular matrix, and increasing cell migration^20, 37, 42, 62^. Hypoxia also upregulates genes which facilitate tumor cell intravasation, survival in the blood stream, extravasation and colonization at distant organs^42^. Consistent with processes associated with hypoxia response, we previously observed angiogenesis defects and a decrease in circulating tumor cells suggesting intravasation defects or an inability for metastatic cells to resist anoikis^29^. Our prior work had shown that these vascular defects were associated with a significant reduction in the major angiogenesis signaling molecule, Vegfa. This study expands on this finding and adds to the growing body of research that has demonstrated E2F1 as a key regulator of angiogenesis^14, 16, 18, 55^; suggesting E2F1 regulates many genes within the hypoxia response program to potentially coordinate the development of tumor vasculature.

In addition to hypoxia response, E2F1^−/−^ tumors had significantly lower expression of genes previously associated with metastasis. *In vitro* studies have demonstrated that Areg^26^, Tead1^81^, Coro1C^69^, Lama5^22^, Tgm2^45^ and Fgf 7^79^ are involved in cell migration and invasion features of tumor cells. The reduced expression of these genes involved in invasion phenotypes may provide additional mechanistic information to explain our previous finding that E2F1^−/−^ tumors had possible invasion/intravasation problems indicated by a reduction in circulating tumor cells^30^.

The data presented here also demonstrated a role for Fgf13 in metastasis. Fgf13 is a nonsecretory protein of the FGF family^75^ and we show that loss of Fgf13 in tumor cells blocked colonization of the lungs in a tail vein injection setting. In order to investigate the mechanism for this block of metastatic capacity, we identified that genes that co-vary with Fgf13 using WGCNA. Amongst these genes, we noted several genes associated with cell motility were in an interaction network with FGF13. This led us to predict that FGF13 may function in cell migration. We confirmed this prediction using a scratch assay where both Fgf13 knockout clones demonstrated impaired cell migration. Interestingly, the role of Fgf13 in neuron development also highlights its role in cell migration while detailing a mechanism of Fgf13 induced microtubule stabilization^75^. Within the Fgf13 covariate network, we also observe many genes consistent with microtubule stabilization and neuron development (Table 1). This agreement between the two studies might suggest conserved mechanisms for cell migration in tumor cells and in neurons. Further, drawing from our study and that of Wu et al.^75^ predicts that the possible function of Fgf13 in metastasis may be attributed to a function in cell migration via microtubule stabilization.

As a whole, this study shows that the metastatic defects associated with E2F1 loss could be due to an inability to properly initiate a gene expression response to hypoxia. Importantly, many of the genes downregulated in E2F1^−/−^ tumors associated with hypoxia response have been shown as regulators breast cancer metastasis. We identified that the E2F1 target gene Fgf13 controls pulmonary metastasis, potentially through a cell migration mechanism; possibly providing a means for E2F1 to regulate cell migration. Collectively, this study furthers our initial characterization of E2F1’s regulation of metastasis by identifying the metastasis associated gene expression response to E2F1 loss.

## METHODS

### RNA AND MICROARRAY

Preparation of RNA samples from flash frozen tumors was done using the Qiagen RNeasy kit after roto-stator homogenization. RNA from 17 Myc induced tumors was submitted to the Michigan State University Genomics Core facility for gene expression analysis using Mouse 430A 2.0 Affymetrix arrays.

### GENE EXPRESSION ANALYSIS

Raw intensity .CEL files were processed and RMA normalized using Affymetrix Expression Console. Gene expression data is deposited on the Gene Expression Omnibus under the accession number GSE104397. Unsupervised class discovery was done using Consensus Cluster Plus^73^. For class discovery, we mapped Affymetrix probes to their gene symbol using the platform table deposited on the Gene Expression Omnibus (GPL8321). Given the presence of multiple probes for single genes, we collapsed duplicate genes to their mean expression in each sample. This approach reduced our dataset of 22690 probes down to 12847 gene symbols. Next, we median centered the gene expression values across the samples in the dataset. Finally, we filtered genes using a requirement of a standard deviation greater than 0.5. This resulted in 1,303 genes used for class discovery and consensus clustering. We used 1,000 iterations and 90% item (sample) resampling to evaluate 2-10 potential clusters (groups/classes). Consensus cumulative distribution function (CDF) plots are also generated using Consensus Cluster Plus^73^. Silhouette width^58^ was used to assess validity of each cluster using the R package ‘Cluster’^52^. Pathway activation was predicted according to previous studies^6, 21, 28^. Single sample gene set enrichment analysis was done for Hallmark^41^ gene sets using the ssGSEAProjection module on Broad Institute’s Gene Pattern^57^. Significance analysis of microarrays^67^ was used to compare E2F^WT/WT^ and E2F1^−/−^ tumors in a fold change analysis. Direct E2F1 target genes were identified using ChIP-base^77^ and data from a previous ChIP-seq experiment^5^. Kaplan-Meier plots were generated using using KMPLOT.com to query human breast cancer expression and clinical data^64^. Significant overlaps with previously established gene sets were detected using the molecular signatures database^41^. Interaction networks were assembled using www.string-db.org^19^. Weighted correlation network analysis was implemented according to published protocols^38^ and using a gene significance score threshold of 0.6 to select genes for further analysis.

### CELL CULTURE

MMTV-PyMT 419 cells were a gift from Dr. Stuart Sell and Dr. Ian Guessand have been previously characterized^44^. All tumor cells were cultured in Dulbecco’s Modified Eagle’s Medium, 3.7 g/L of NaHCO3, 3.5 g/L d-glucose, 5ug/mL insulin, 1ug/mL hydrocortisone, 5ng/mL Egf, 35ug/mL BPE, 50ug/mL gentamicin, 1X Antibiotic/Antimycotic, and 10% fetal bovine serum. Media was set to a pH of 7.4.

### CRISPR

Sequence for Fgf13 was obtained from the UCSC genome browser^36^. Guide sequences were predicted by submitting exon (using only those that were common across all Fgf13 isoforms) sequence using the CRISPR design tool at: http://crispr.mit.edu/. Oligos for guide sequence assembly were designed by adding a ‘G’ followed by ‘CACC’ at the 5’ end of the guide sequence. For the complementary DNA to the guide, add ‘CAAA’ to the 5’ end. Oligonucleotide sequences are as follows:

Fgf13 5’: CACCGTCAGCAGCAATCCGGCCGA

Fgf13 3’: AAACTCGGCCGGATTGCTGCTGACC

Oligonucleotides for guide sequence assembly were ordered from integrated DNA technologies https://www.idtdna.com/site. Oligos were diluted to a concentration of 100uM in water. To anneal the oligonucleotides 5 uL of the forward and 5uL of the reverse oligo are incubated in 10uL of 2X annealing buffer (10 mM Tris, pH 7.5–8.0, 50 mM NaCl, 1 mM EDTA) at 95 degrees Celsius for 4 minutes, and then cooled to room temperature. The annealed oligonucleotides were inserted into the PX458 vector from Addgene (#48138). Confirmation of the cloned guide sequence was done using Sanger sequencing.

PyMT 419 cells were transfected according to a previously described protocol^40^. To select clones, GFP positive cells were sorted into 96 well plates using fluorescence activated cell sorting. Knockout clones were screened for by PCR amplifying a ~300 bp amplicon with the PAM sequence centrally located within the PCR product. PCR products were resolved on a 3% agarose gel, extracted with the QIAquick Gel Extraction Kit, and were submitted for sanger sequencing.

PCR Amplification Primers:

Fgf13 5’: 5’-TGTTCTAACTTCCAGAAAGGCATA-3’

Fgf13 3’: 5’-CAGTGGTTTGGGCAGAAAAT-3’

For sequencing, a nested primer with the following sequence was used.

Fgf13 5’: 5’-CACACCCATATAAGTATTGACTTTCA-3’

Knockout and add-back were confirmed by western blot according to published methods^27^. For western blots, we used a polyclonal Fgf13 antibody (Invitrogen PA527302) and for beta-actin we used a monoclonal antibody (Cell Signaling #4970). The Fgf13 ORF was purchased from GenScript in the pcDNA3.1 vector and were subsequently digested with EcoRI and XhoI cloned into the EcoRI and XhoI sites of pLXSN. Retrovirus was packaged in HEK-293GPG as previously described (90). Briefly, a confluent 10-cm plate of HEK-293GPG cells was transfected with 5 µg of pLXSN-mADM or pLXSN-mFGF13 and virus containing media was subsequently harvested. Subconfluent 10 cm plates (~20%confluence) of 419 cells were infected with 1 mL of non-concentrated viral supernatant and infected cells were selected with 300 µg/mL G418.

### IN VITRO ASSAYS

To measure cell migration, wound healing assays were performed in the presence of 2ug/mL Mitomycin C using standard methods^60^. Photomicrographs were taken at 0 hour and 18 hours.

### IN VIVO ASSAYS

Animal protocols used for this study were approved by Michigan State University IACUC Committee and conducted according to national and institutional guidelines. All mice were in the FVB background. For tail vein injection, MMTV-Cre control mice were used to avoid immune response to the middle T antigen^17, 43, 56^. For control cells and each knockout clone, 50,000 cells were injected into the bloodstream via the tail vein. After 21 days, mice were euthanized. Lungs were resected for H&E staining to detect metastases. To calculate significance, unpaired T-tests were done in graphpad prism.

## ADDITIONAL INFORMATION

### Author Contributions

DPH and ERA conceived and designed the work. DPH and ERA wrote the manuscript. DPH performed the Crispr gene targeting, RNA isolation, cell counts, migration assays, gene expression analysis, tail-vain injection studies. JPR assisted with Crispr vector construction and mouse work. SAM assisted with tissue culture and knockout screening and sequencing. BCM provided in vitro analysis. AM provided tissue culture and DNA isolation support. KAG and KMC provided reagents and experimental guidance. All authors reviewed the manuscript.

## Acknowledgments

We thank Dr. Stuart Sell and Dr. Ian Guess for providing their PyMT 419 cells for this study.

