## Supplemental Figures for "E2F1 Drives Breast Cancer Metastasis by Regulating the Target Gene FGF13 and Altering Cell Migration"

**Supplemental Figure Legends**

Figure S1- Hiearchical clustering using genes for intrinsic subtype.

(1A) Hierarchical clustering of MMTV-PyMT tumors following BFRM [1] batch correction with MMTV-Neu and MMMT-Myc tumors using a published intrinsic gene list[2]. The dendrogram illustrates relationships amongst samples coming from each model, as indicated by green (PyMT), blue (Neu), or black bars (Myc). Besides the heatmap purple bars indicate the position of gene clusters highlighted in panel B. A total of 1,420 genes were clustered and expression levels are illustrated by the color bar. (B) Detailed analysis of for clustering shown in panel A. Genotypes and tumor histology is shown by the color coded bars for each model as described. Highlighted gene clusters are as follows: (i) basal-like, (ii) luminal-like, (iii) proliferation, and (iv) claudin-low.

Figure S2 – Analysis of proliferation and tumor growth rate in MMTV-PyMT histological subsets.

(A) Relative expression level of a published gene expression signature for proliferation [3] across the various histological types of MMTV-PyMT tumors. Scores are depicted as the median expression of the gene set in each tumor. (B) Average tumor growth rate by histology as calculated by the number of days from tumor palpation until the primary tumor reaches a volume of 2000 $mm^3$.

Figure S3- Expression level of genes with demonstrated metastatic function in previous *in vivo* studies.

The RMA normalized median centered expression value for genes with demonstrated in vivo mediation of metastasis is shown in E2F WT and E2F1 KO tumors for (A) Vegfa [4], (B) Hbegf [5], (C) Hspb1[6], (D)Flt1[7], (E) L1cam [8], and (F) Plaur [9]. Two-tailed p-values are as depicted above the boxplots resulting from an unpaired t-test .

Figure S4- Expression level of genes with demonstrated metastatic function in previous *in vitro* studies.

The RMA normalized median centered expression value for genes with demonstrated *in vitro* mediation of metastasis is shown in E2F WT and E2F1 KO tumors for (A) Areg [10], (B) Tead1 [11], (C) Coro1C [12], (D) Lama5 [13], (E) Tgm2 [14], and (F) Fgf 7 [15] . Two-tailed p-values are as depicted above the boxplots resulting from an unpaired t-test .

### Supplemental Figure Legend References


1 Carvalho, C. M. *et al.* High-dimensional sparse factor modeling: applications in gene expression genomics. *Journal of the American Statistical Association* **103**, 1438-1456 (2008).

2 Pfefferle, A. D. *et al.* Transcriptomic classification of genetically engineered mouse models of breast cancer identifies human subtype counterparts. *Genome Biol* **14**, R125 (2013).

3 Fan, C. *et al.* Building prognostic models for breast cancer patients using clinical variables and hundreds of gene expression signatures. *BMC medical genomics* **4**, 1 (2011).

4 Schoeffner, D. J. *et al.* VEGF contributes to mammary tumor growth in transgenic mice through paracrine and autocrine mechanisms. *Laboratory investigation* **85**, 608-623 (2005).

5 Zhou, Z. *et al.* Autocrine HBEGF expression promotes breast cancer intravasation, metastasis and macrophage-independent invasion in vivo. *Oncogene* **33**, 3784-3793 (2014).

6 Gibert, B. *et al.* Targeting heat shock protein 27 (HspB1) interferes with bone metastasis and tumour formation in vivo. *British journal of cancer* **107**, 63-70 (2012).

7 Taylor, A. P. & Goldenberg, D. M. Role of placenta growth factor in malignancy and evidence that an antagonistic PlGF/Flt-1 peptide inhibits the growth and metastasis of human breast cancer xenografts. *Molecular cancer therapeutics* **6**, 524-531 (2007).

8 Zhang, H. *et al.* HIF-1-dependent expression of angiopoietin-like 4 and L1CAM mediates vascular metastasis of hypoxic breast cancer cells to the lungs. *Oncogene* **31**, 1757-1770 (2012).

9 Xing, R. H. & Rabbani, S. A. Overexpression of urokinase receptor in breast cancer cells results in increased tumor invasion, growth and metastasis. *International journal of cancer* **67**, 423-429 (1996).

10 Higginbotham, J. N. *et al.* Amphiregulin exosomes increase cancer cell invasion. *Current Biology* **21**, 779-786 (2011).

11 Zhang, H. *et al.* TEAD transcription factors mediate the function of TAZ in cell growth and epithelial-mesenchymal transition. *Journal of biological chemistry* **284**, 13355-13362 (2009).

12 Wang, J. *et al.* miR-206 inhibits cell migration through direct targeting of the actin-binding protein Coronin 1C in triple-negative breast cancer. *Molecular oncology* **8**, 1690-1702 (2014).

13 Giannelli, G., Falk-Marzillier, J., Schiraldi, O., Stetler-Stevenson, W. G. & Quaranta, V. Induction of cell migration by matrix metalloprotease-2 cleavage of laminin-5. *Science* **277**, 225-228 (1997).

14 Mangala, L., Fok, J., Zorrilla-Calancha, I., Verma, A. & Mehta, K. Tissue transglutaminase expression promotes cell attachment, invasion and survival in breast cancer cells. *Oncogene* **26**, 2459-2470 (2007).

15 Zang, X. P. & Pento, J. T. Keratinocyte growth factor-induced motility of breast cancer cells. *Clinical & experimental metastasis* **18**, 573-580 (2000).

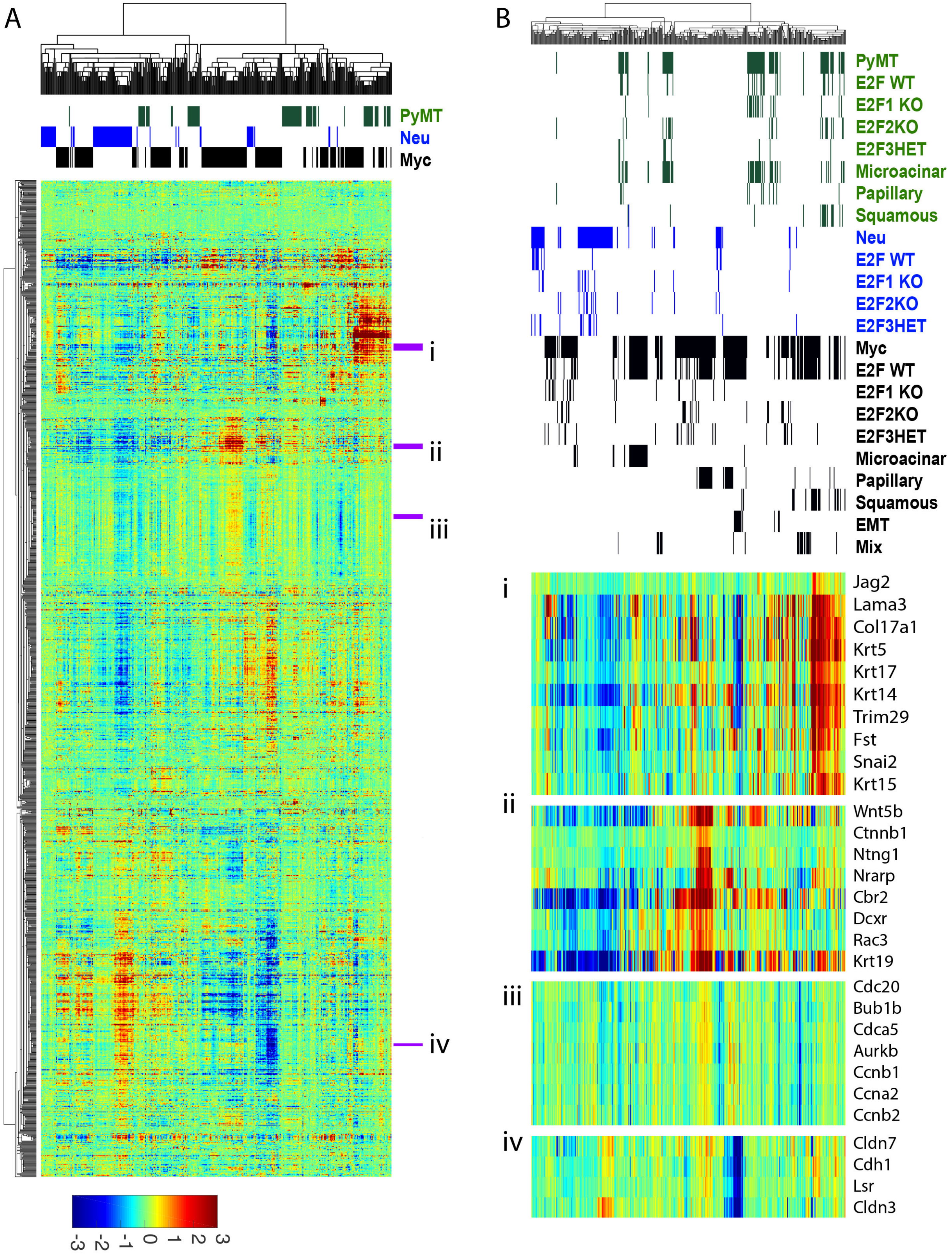
A
PyMT
Neu
Myc
i
ii
iii
iv
-3 -2 -1 0 1 2 3
B
PyMT
E2F WT
E2F1 KO
E2F2KO
E2F3HET
Microacinar
Papillary
Squamous
Neu
E2F WT
E2F1 KO
E2F2KO
E2F3HET
Myc
E2F WT
E2F1 KO
E2F2KO
E2F3HET
Microacinar
Papillary
Squamous
EMT
Mix
i
Jag2
Lama3
Col17a1
Krt5
Krt17
Krt14
Trim29
Fst
Snai2
Krt15
ii
Wnt5b
Ctnnb1
Ntng1
Nrarp
Cbr2
Dcxr
Rac3
Krt19
iii
Cdc20
Bub1b
Cdca5
Aurkb
Ccnb1
Ccna2
Ccnb2
iv
Cldn7
Cdh1
Lsr
Cldn3

A

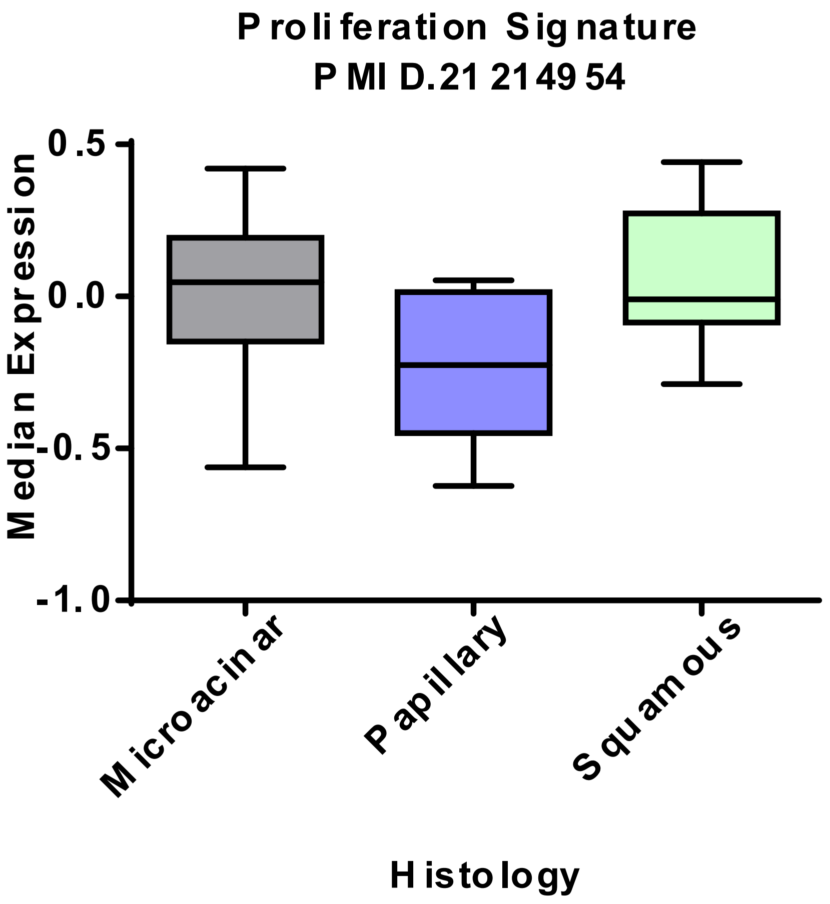

Proliferation Signature
PMID.21214954
Median Expression
0.5
0.0
-0.5
-1.0
Microacinar
Papillary
Squamous
Histology


B

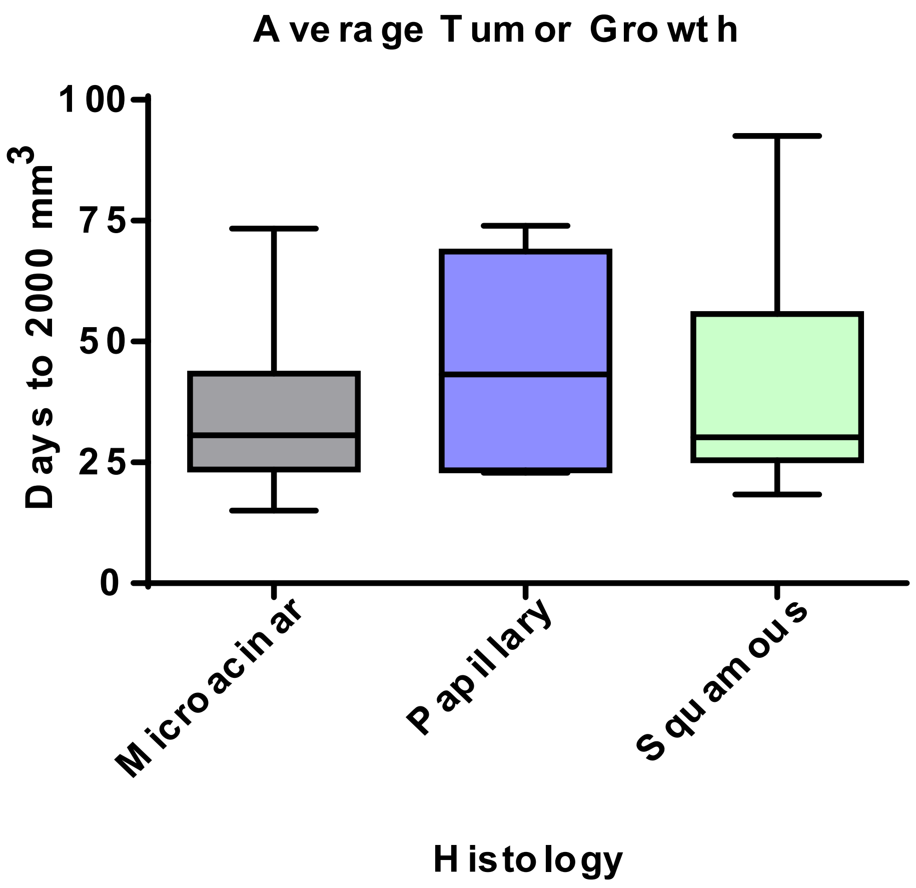

Average Tumor Growth
Days to 2000 mm3
100
75
50
25
0
Microacinar
Papillary
Squamous
Histology

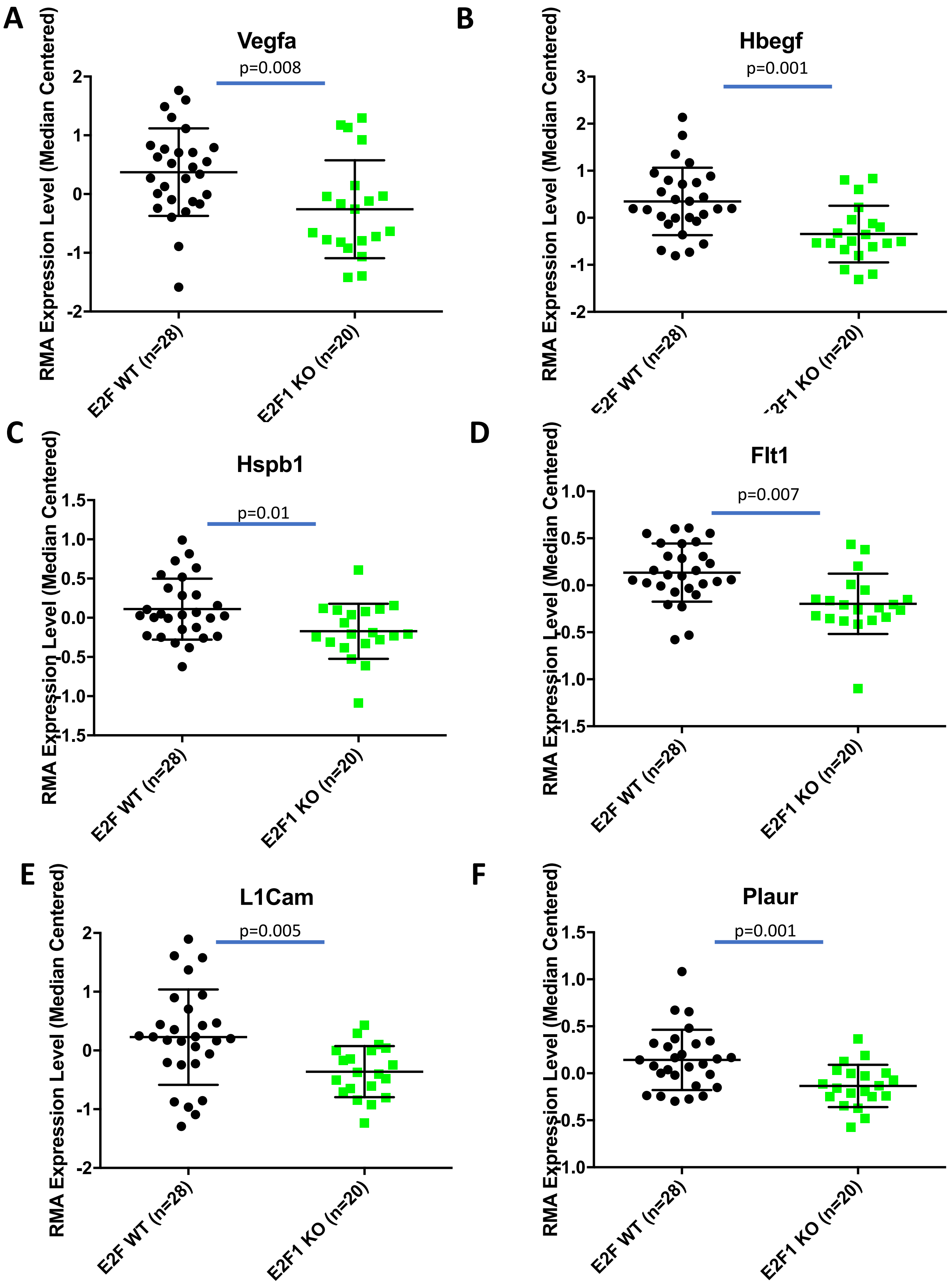
A
Vegfa
p=0.008
RMA Expression Level (Median Centered)
E2F WT (n=28)
E2F1 KO (n=20)
B
Hbegf
p=0.001
C
Hspb1
p=0.01
D
Flt1
p=0.007
E
L1Cam
p=0.005
F
Plaur
p=0.001

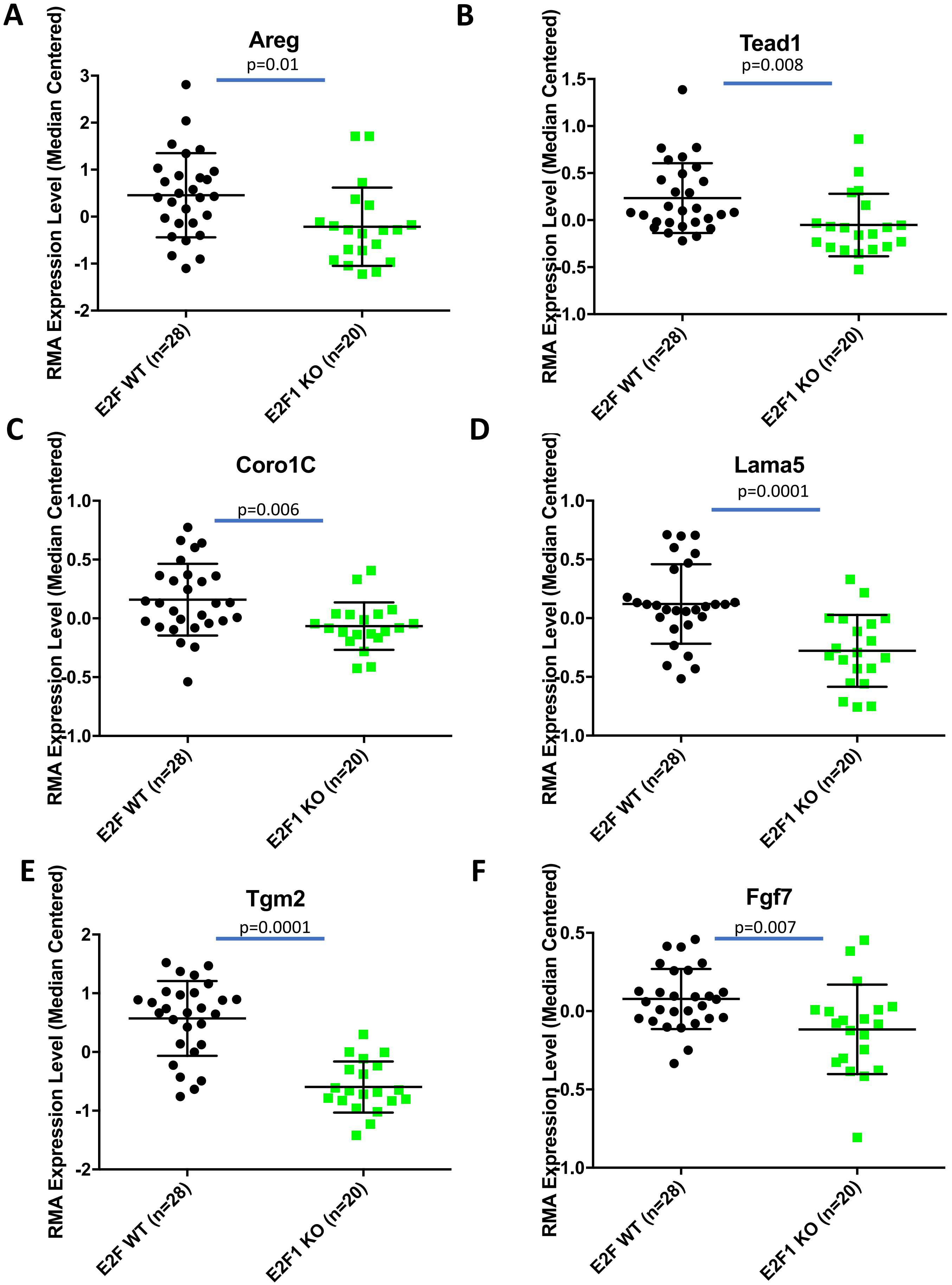
A
Areg
p=0.01
B
Tead1
p=0.008
C
Coro1C
p=0.006
D
Lama5
p=0.0001
E
Tgm2
p=0.0001
F
Fgf7
p=0.007
RMA Expression Level (Median Centered)
E2F WT (n=28)
E2F1 KO (n=20)
